# Mechanistic insights into transport models of the sphingolipid transport protein, Spinster homolog 2 (Spns2), using MD simulations

**DOI:** 10.1101/2024.03.10.584301

**Authors:** Amanda K. Sharp, Marion Q. LoPresti, Anne M. Brown

## Abstract

Sphingosine-1-phosphate (S1P) is a sphingolipid signaling molecule that when elevated results in multiple disease states including metastatic cancers. Modulating the extracellular concentrations of S1P has been an evolving strategy in drug development for metastatic cancers due to its role in angiogenesis and cell migration. Research has shown that Spns2, the S1P transport protein, is an important microenvironment regulatory gene in metastatic lung cancer colonization and has demonstrated that Spns2 inhibition is a powerful suppressor of metastatic cancers. Spns2 transports and regulates cellular levels of S1P but has unresolved aspects related to mechanism of transport. Here, molecular modeling strategies including, homology modeling and molecular dynamics (MD) simulations, were used to determine structural mechanisms of action related to S1P transport and exploitable for inhibition. Results indicate Spns2 contains a unique salt-bridge network essential for structural stability that is disrupted by the R119A mutation. Additionally, we observe that Spns2 follows a rocker-switch transport model and that S1P translocation is initialized by interacting with residues such as Thr216, Arg227, and Met230. This work provides initial insight into structural morphologies sampled by Spns2, the role of a complex salt bridge network, and residues engaged in structural state transition that can be targeted with inhibitors to control extracellular concentrations of S1P.

## Introduction

Metastatic cancer, or stage 4 cancer, occurs when the cancer cells spread from the originating cancer site to other tissues and organs. This process can lead to deterioration to adjacent organs, causing loss of organ function, leading to death. 90% of cancer-related deaths are from metastatic cancer with no success in improving this statistic in the last 50 years.(*1, 2*) Microenvironment regulation is essential for malignant tumors to metastasize, and specific genes have been shown to enable routes for metastatic motility in breast, prostate, and lung cancers.(*3*) A microenvironment that supports angiogenesis is essential for cancer cells to initiate motility and infiltrate other organs.(*3*) Identifying avenues of metastasis-enabling microenvironments that can be exploited for therapeutic development can provide alternative routes for cancer drug discovery opportunities. S1P is a sphingolipid signaling molecule has been implicated in numerous cellular processes that support cancer survival, metastasis, and chemoresistance. S1P is involved in the ceramide synthase pathway and is tightly regulated with ceramide to promote cellular homeostasis.(*4*) An imbalance in the ceramide/S1P ratio has been shown to promote diseases including various cancer types.(*5, 6*) S1P acts via a paracrine and autocrine action to stimulate inflammatory cell trafficking, angiogenesis, cell migration, and cellular proliferation, providing a favorable microenvironment for cancer metastasis.(*7–9*) Elevated cellular concentrations of S1P have been linked to dysregulation of cellular processes and increase risk of disease development.(*10, 11*) Specifically, extracellular S1P has been shown to stimulate cellular angiogenesis,(*12*) providing a route for tumor or cancer cell migration. For these reasons, exploring multiple strategies to lower cellular concentrations of S1P has become an attractive research strategy for therapeutic development of metastatic cancers.

S1P is shuttled out of the cell via the Spns2 transport protein. Spns2 is an ATP-independent transport protein from the Major Facilitator Superfamily (MFS). MFS are a large family of secondary transporters with over 75 different subfamilies that share a common 12-transmembrane helical domain comprised of two 6-transmebrane bundles.(*13, 14*) MFS members are secondary active transport proteins, which act by either symport or antiport process, to provide an electrochemical gradient to support solute carrier transport.(*15*) MFS members also follow three distinct models dependent on orientation of their helical bundles for transporting substrates across a membrane: rocker-switch, rocking-bundle, or elevator models.(*15*) Spns2 is a member of the endosomal spinster family within the MFS 2.A.1.49 superfamily,(*16*) with three family members, Spns1, Spns2, and Spns3, all of which have little known structural information including secondary active transport process, substrate transport mechanism, or overall function.

Spns2 has become an emerging protein target of interest for pharmaceutical development for metastatic cancers due to its role in transporting S1P, leading to elevated circulating concentrations of S1P and extracellular signaling, as well as promising results with genetic studies emphasizing its importance in metastatic cancer.(*11*) Additionally, sequence analysis of Spns2 has shown that the R119 in Spns2 appears to be essential for S1P secretion and the R119A mutant significantly decreases extracellular concentrations of S1P.(*17*) Spns2 also contains an arginine-rich cytofacial domain that may play a role in the recruitment of S1P for transport. Recently, Spns2 crystal structures have been resolved demonstrating three distinct states: inward-facing (PDB ID: 7YUF), outward-facing partially occluded (PDB ID: 8EX7 and 8EX8), and outward-facing (PDB ID: 8EX5). Additional structures have been resolved with inhibitor 16d (PDB ID: 8G92, 8JHR, 8KAE) and substrate S1P (PDB ID: 8EX4, 8EX6, 8JHQ, 7YUB).(*18–20*) The inward-facing state is defined by a cytofacial solvent accessible cavity, whereas the outward-facing state is solvent accessible on the exofacial side. The outward-facing partially occluded state is identified during the transition from the inward-facing to outward facing state.(*18*)

Previous research has focused on exploring the *Hyphomonas neptunium* homolog of Spns2 to provide structural insight and has identified a proton-mediated secondary transport mechanism with this homolog, but this limits our understanding into the human function as its similarity is low (28%). Additionally, researchers have identified potent Spns2 inhibitors but there is still minimal understanding into its structure and functional mechanisms.(*21*) To better target S1P extracellular concentrations via Spns2, a better understanding of dynamics-based structural functionality is necessary.

Understanding the mechanism of S1P transport is essential for the development of therapeutic agents. To explore this mechanism, we employed homology modeling coupled with molecular dynamics (MD) simulations of Spns2 in a membrane environment to identify the changes in secondary active transport influence by the R119A mutation. Identifying the structural influences of S1P mechanistic transport of Spns2 in a membrane environment that can lead to an improved understanding of strategies to inhibit S1P translocation, providing additional routes to control extracellular levels of S1P. We identified that the Spns2 follows a rocker-switch transport model, there is an inward-facing occluded state not defined by resolved structures, Spns2 contains an inter-bundle salt-bridge network essential for overall structural stability that is disrupted by the R119A mutation, and that S1P translocation is initialized by interacting with residues such as Thr216, Arg227, and Met230. This work delivers a unique insight into Spns2 structural mechanisms and provides an improved understanding on strategies to inhibit S1P translocation, offering additional routes to control extracellular levels of S1P.

## Materials and Methods

### Homology Modeling

The sequence of Spns2 was obtained from Genbank(*22*) (accession: NP_001118230.1). The Iterative Threading ASSEmbly Refinement (I-TASSER) server was utilized to employ homology modeling of the Spns2_WT_ and Spns2_R119A_ structures using residues 100-549 and the template of the *Hyphomonas neptunium* homolog of Spns2 (PDB 6e8j).(*23*) Spns2 and HnSpns2 sequences displayed a 28% similarity, 18% identity, and 36% homology (**Figure S1**). Energy minimization was performed on the highest ranked model based on highest confidence score (C-score) using the protein preparation wizard and the OPLS3e forcefield within Maestro 12.3 as a part of the Schrödinger Release 2020-1 (5–7). To structurally validate the models, each model was analyzed using Ramachandran plots(*24*), and ProSA(*25*) (**Figure S2-3**).

### Molecular Dynamics Simulations (MDS)

Molecular dynamics (MD) simulations were employed to probe the dynamic profile of Spns2. Four total systems were generated to be ran through MD simulations: Spns2_WT_ apo, Spns2_R119A_ apo, Spns2_WT_ with S1P, and Spns2_R119A_ with S1P. To build the MD simulation membrane environments, the CHARMM-GUI membrane builder was used(*26*). The extracellular membrane leaflet was constructed to replicate a general plasma membrane with 219 cholesterol, 280 1-palmitoyl-2-oleoyl-*sn*-phosphatidylcholine (POPC), 42 1-palmitoyl-2-oleoyl-*sn*-phosphatidylethanolamine (POPE), and 153 palmitoylsphingomyelin (PSM) molecules while the intracellular membrane leaflet contained 196 cholesterol, 182 1-palmitoyl-2-oleoyl-*sn*-phosphatidylethanolamine (POPE), 70 3-palmitoyl-2-oleoyl-D-glycero-1-phosphatidylserine (POPS), 42 1-palmitoyl-2-oleoyl-inolsitol (POPI), 70 PSM, and 140 POPC molecules.(*27*) Each system was solvated in a mixed cationic environment with 75 mM KCl and 75 mM NaCl net neutral system with the TIP3P water model.(*28*) Random placements of S1P were generated using the GROMACS insert-molecules features using a de-solvated Spns2 membrane system and selected for placement near the cytofacial domain of Spns2. S1P systems were re-solvated using GROMACS. Constructed systems were simulated using GROMACS v. 2020.3 software(*29*) using the CHARMM36m forcefield(*30*). Energy minimization used the steepest descent method along with a tolerance of 1000 kJ mol^-1^ nm^-1^. Each system generated replicates starting at the first equilibration step. Apo systems each were generated with four replicates at the first equilibration step and S1P systems with six replicates. Equilibration was performed at 310 K temperature and 1 bar pressure and used a six-step process. The first step applied random velocities and used the Berendsen thermostat temperature coupling method(*31*). The second step was an extension of the first step, employing the same parameters. The third step applied expanded by using a semi-isotropic coupling type(*31*). Steps four, five, and six used the same parameters as the third step with a stepwise release of position restraints on the system. MD production applied the Verlet cutoff-scheme used for long distance cutoffs with 1.2 Å distance for neighbor searching,(*32*) Linear constraint solver constraint-algorithm(*33*), the Nose-Hoover temperature coupling algorithm(*34, 35*), and the smooth particle-mesh Ewald (PME) applied to hydrogen-bonds(*36, 37*). MD was run with a 310 K temperature and 1 bar pressure environment. Each system replicate was simulated for a total of 1000 ns.

### MDS Analysis

Analysis was performed MD simulations using GROMACS(*29*) analysis suite and in-house produced analysis scripts. Apo systems used an analysis timeframe over the last 200 ns of simulated structures based off root-mean-square deviation (RMSD) calculations to identify system stability. Clustering of dominant morphologies were performed on the last 200 ns on full concatenated systems and individual replicate structures with a 0.2-nm cutoff using the gromos algorithm. Distance calculations were performed using the GROMACS mindist and distance calculation features. Angle calculations were performed using the MDtraj python package and an in-house produced python script using the Z and Y coordinates after inspecting the 3D positionings using matplotlib. Ion distances to the membrane normal were performed using the distance GROMACS feature and using the z-axis output. Residue-ion interactions were performed using an in-house produced python script. Principal component analysis was performed using GROMACS covar and anaeig features to compare phase-space between systems. In-house analysis scripts, starting simulation structures, and topology data can be found on our public Open Science Framework page (https://osf.io/82n73/).

## Results and Discussion

Cancer metastasis is the cause of 90% of cancer-related deaths due to its position in disease progression.(*1, 2*) Identifying avenues to suppress cancer into progressing and infiltrating other tissue can help improve cancer survival statistics overall. Sphingosine-1-phosphate (S1P) has become a focus of study in the metastatic cancer research field as it has been identified to support a metastasis-enabling environment, driving cell motility and disease progression.(*17*) As Spns2 is the dominant transport protein of S1P, it serves as a strategic avenue for controlling S1P transport. Here, molecular modeling and atomistic MD simulations were used to explore the morphological states sampled of Spns2. This work found that Spns2 likely follows a rocker-switch transport model, that there is an inward-facing occluded state not identified in currently resolved structures, that the R119A mutation impacts helical plasticity in the gating and cavity helices, and that R119A impacts a salt-bridge network essential in protein stability.

Homology models of Spns2_WT_ and Spns2_R119A_ were generated using the *Hyphomonas neptunium* homolog of Spns2 (PDB 6e8j),(*23*) where sequences displayed a 28% similarity, 18% identity, and 36% homology (**Figure S1-S3**). Four systems were constructed for MD simulation input: Spns2_WT_ apo, Spns2_R119A_ apo, Spns2_WT_ with S1P, and Spns2_R119A_ with S1P using the CHARMM36 forcefield(*30*). To identify system stability, root mean square deviation (RMSD) and RMSD clustering was used, demonstrating that the last 200 ns of simulation were adequate for sampling coverage to understand an unchanging state of the structure (**Figure S4**). Clustering of the concatenated systems over the last 200 ns were performed to identify dominant morphologies (**Figure S5-6**), demonstrating well preserved helical bundles throughout the different systems.

### Spns2 follows a rocker-switch transport model and R119A influences helical distance changes within the gating and cavity helices

MFS members follow three distinct models dependent on orientation of their helical bundles for transporting substrates across a membrane: rocker-switch, rocking-bundle, or elevator models.(*15*) Spns2 has been hypothesized to follow a rocker-switch model based on MD simulations of the *Hyphomonas neptunium* homolog of Spns2 (HnSpns2).(*38*) To further explore this hypothesis, we analyzed the Spns2 N-terminal and C-terminal helical bundles (**Figure 1A**) for their angles between each other and relative to the membrane normal (**Figure 1B**). The angles between the two helical bundles were highly flexible in the Spns2_WT_ system, demonstrating helical bundle plasticity with an average of 37° ± 6 over the last 250 ns. Spns2_R119A_ system was observed to have a larger angle between the two helical bundles with more bundle rigidity, with an average of 57° ± 5. Assessing the angles between the helical bundles against the membrane normal, we observed that both helical bundles maintained a tilted orientation within the membrane and similar angles, showing the N-terminal mean of 52° ± 1 and C-terminal mean of 51° ± 1 (**Figure 1C**). The similarity of the angles against the membrane normal is consistent of a rocker-switch mechanism, whereas both rocking-bundle and elevator models would demonstrate a singular bundle closer to a 90° angle.(*15*) We also analyzed the Spns2_R119A_ system to detect if the single-point mutation obstructs the substrate-transport model and minimal differences were observed (N-terminal mean of 52° ± 1 and C-terminal mean of 48°± 1). This data further supports that Spns2 follows a rocker-switch model for substrate transport, previously identified from work analyzing HnSpns2. Additionally, these observations support that the R119A mutation impacts the helical bundle angle distances, potentially impacting substrate translocation via a disruption of the cytofacial binding interface.

**Figure 1.**
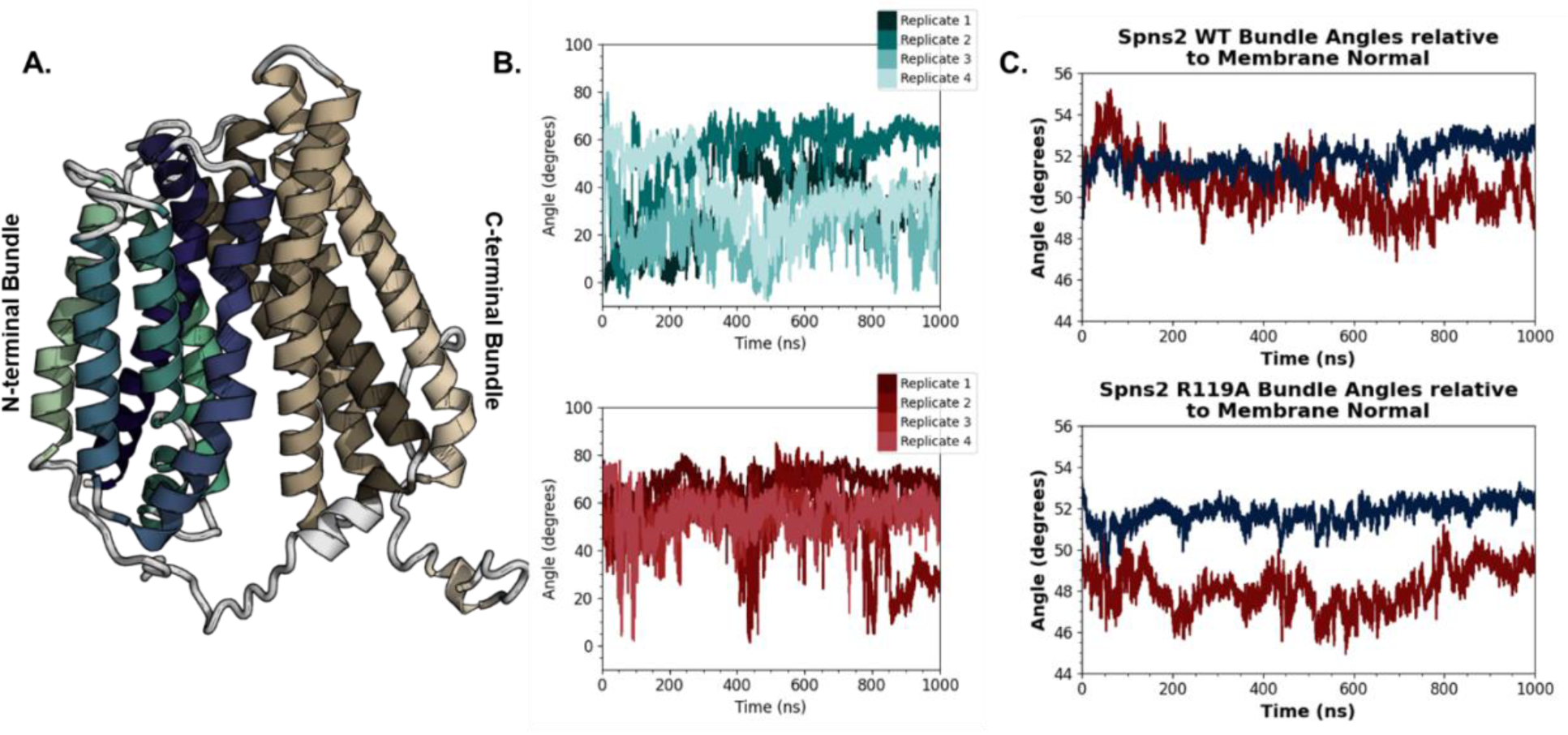
Spns2 follows a rocker-switch transport mechanism. A. Visualization of helical bundles with green and teal helices representing the N-terminal bundle and tan and brown helices representing C-terminal bundle. B. Bundle-bundle angle calculations. Angle calculations were determined using Cα atoms and were calculated using MDtraj, exporting the Y and Z coordinates. C. Angle calculations of each helical bundle relative to the membrane normal calculated using GROMACS distance.

Research has shown that interactions between helices 2 & 8 (H2-H8) and 5 & 11 (H5-H11) are “gating helices” involved in inter-domain conformational shifts important in substrate translocation (**Figure 2A**).(*38*) These gating helices are centrally located between the two helical bundles. Helix distance frequencies demonstrate a difference between H2-H8 in comparing the Spns2_WT_ and Spns2_R119A_ systems in the cytofacial region (**Figure 2B**), showing that the helical distances are further in the Spns2_WT_ system and that H5-H11 were shown to have differences in distance probabilities (**Figure 2B**). Both systems demonstrate bimodal distribution distances between H5-H11, where H5 is approximately 1-2 nm closer to H11 in the WT system as compared to the R119A system. These results suggest that H2-H8 being further apart in the cytofacial region supports substrate binding, and the R119A mutation may impact the gating function within the protein. Additionally, helices 1, 4, 7 and 10 are considered “cavity helices” which are responsible for substrate binding (**Figure 3A**). Analysis of distance probabilities for those helices on the cytofacial range show most displacement is between H1 with the other cavity helices, with additional differences between H7 and H10 (**Figure 3B**). The R119A mutation, located within H1, appears to directly impact neighboring cavity helices, which could explain the loss of function that has been experimentally determined.

**Figure 2.**
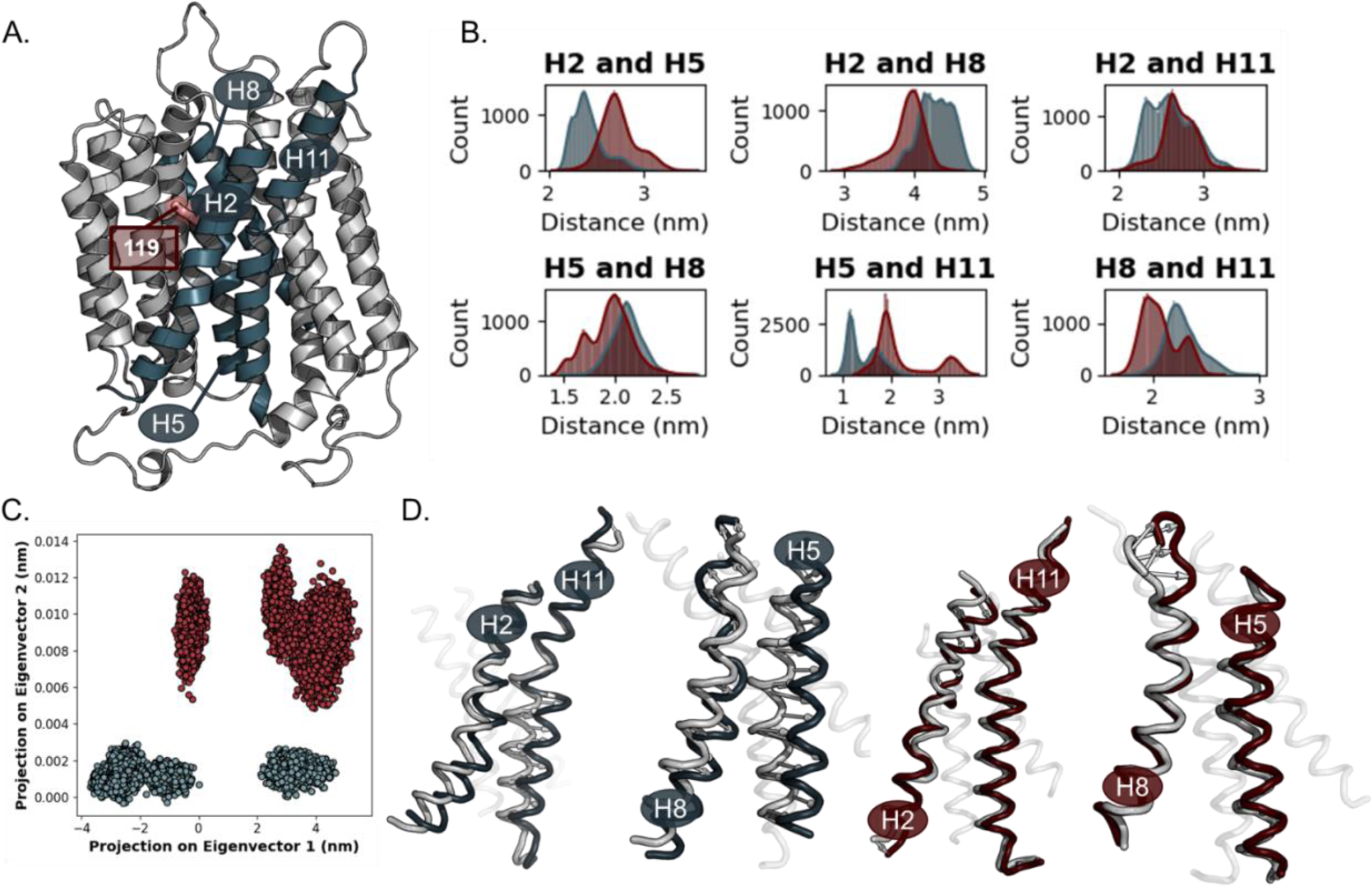
R119A alters gating helical interaction distribution. A. Gating helices highlighted in full structure of Spns2 in teal, with the 119-position colored in red. B. Helical distance distributions in the cytofacial region. Helix distances were calculated using GROMACS mindist feature. C. Principal component analysis of WT and R119A gating helices throughout full simulation to understand phase space differences between the two systems. D. Extreme positions of gating helices of WT (blue) and R119A (red), exported from the GROMACS anaeig feature.

**Figure 3.**
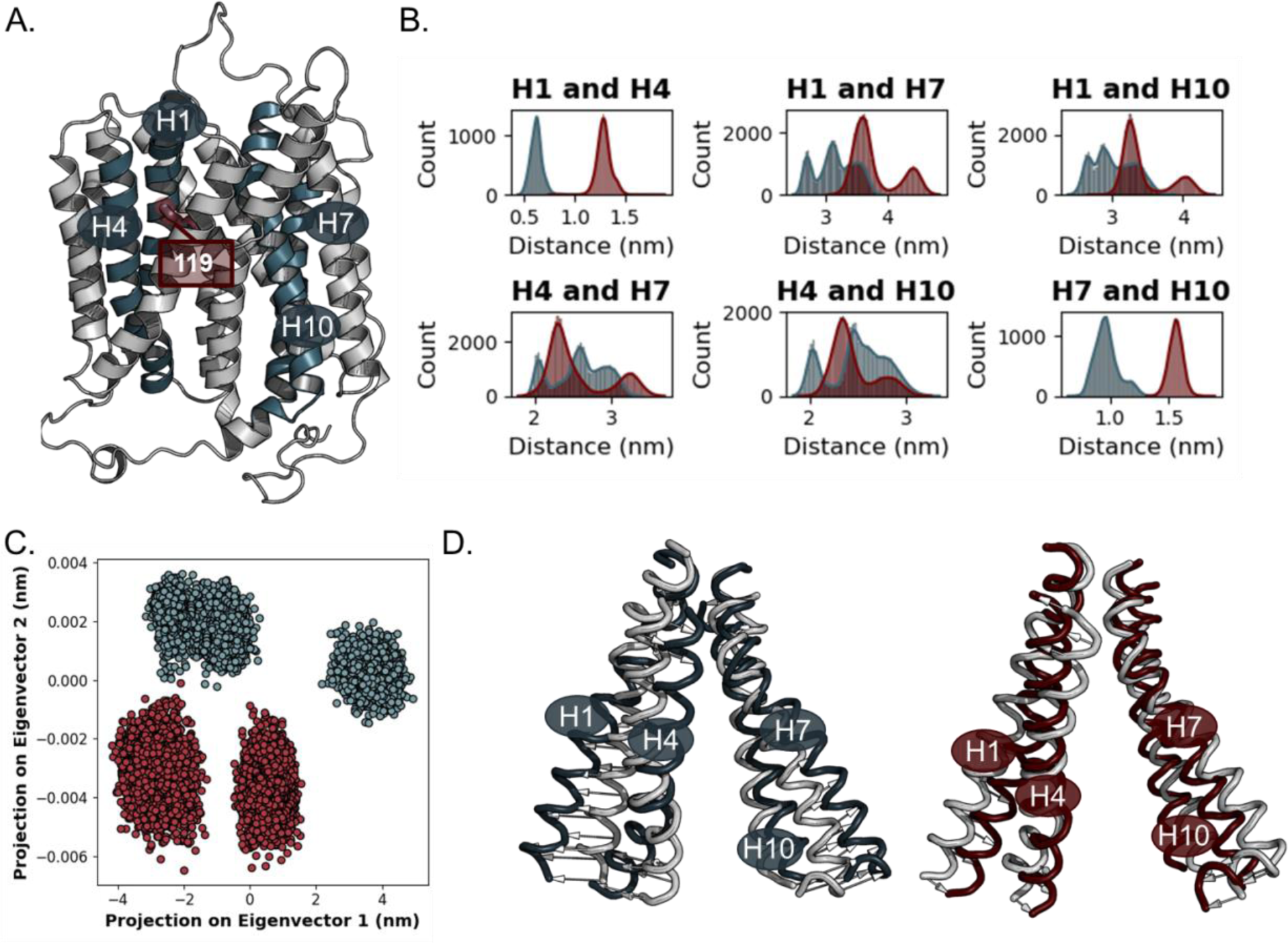
R119A alters cavity helical interaction distribution. A. Cavity helices highlighted in full structure of Spns2 in teal, with the 119-position colored in red. B. Helical distance distributions in the cytofacial region. Helix distances were calculated using GROMACS mindist feature. C. Principal component analysis of WT and R119A gating helices throughout full simulation to understand phase space differences between the two systems. D. Extreme positions of gating helices of WT (blue) and R119A (red), exported from the GROMACS anaeig feature.

These gating helices movements also allow for the classification of certain states. New Spns2 crystal structures show the inward-facing (PDB ID: 7YUF), outward-facing partially occluded (PDB ID: 8EX7 and 8EX8), and outward-facing (PDB ID: 8EX5) structures elucidating those specific states between the transitions, however this analysis proves that there may be an inward-facing occluded state as well. This state is characteristic of the inward-facing state, however, unlike the inward-facing state, TM11 and TM5 have occluded the cytofacial side of the channel. This inward-facing partially occluded state occurs due to the inward movement of an intracellular alpha-helices between TM6 and TM7 pushing TM11 towards TM5 on the cytofacial side effectively occluding the cytofacial pocket. This causes TM10 to move outward, opposite of the direction it would travel to reach the outward-facing state. These movements create two main pockets inside the channel. When exploring potential areas where inhibtiors or other molecules could interact within the channel, there appear to be two binding sites. The first binding site is deeper within the transport channel compared to the second binding site closer to the intracellular side. BS1 mainly interacts with residues from TM7, 8, 10, and 11 including Thr-370, Phe-437, and Asp-472 whereas the second binding site mainly interacts with residues from TM1, 4, 5, and 11 including Asn-112, Pro-215, and Phe-465. Interestingly, BS1 is the same binding site targeted by S1P in crystal structures.

Further analysis of the gating and cavity helices was performed using Principal Component Analysis (PCA). The results demonstrate significant differences in sampled space between the two systems (**Figure 2C-D**, **3C-D**). Inspecting the extreme positions of the gating helices during simulation reveals that helices 5 and 11 orientation appear to be most impacted by A119, where Spns2_WT_ displays more helical plasticity while Spns2_R119A_ had more helical rigidity overall and slightly more flexibility in the extrofacial region (**Figure 2D**). With the cavity helices, similar observations are shown with helices 1, 4, and 7 demonstrating more helical flexibility and scissor-like motion within the cytofacial region of Spns2_WT_ while Spns2_R119A_ shows more helical rigidity (**Figure 3D**). Local changes within the cavity helices have been identified to support the bind and release of substrates and the change in helical plasticity in the region may impact either of those functions.(*15*) Further analysis of why this difference is occurring and if these structural differences impact secondary active transport is necessary.

### R119A disrupts the Spns2 inter-bundle salt-bridge network via alterations to the electrochemical gradient

MFS members are secondary active transport proteins, which act by either symport or antiport process, to provide an electrochemical gradient to support solute carrier transport.(*15*) To analyze the Spns2 secondary active transport model and if secondary active transport is impacted by the R119A mutation, we analyzed the distance between the different ion types to the center of the membrane (**Figure 4A-B**). We observed that in Spns2_WT_ that K^+^ is the dominant interacting ion. Interestingly, Spns2_R119A_ systems displayed interactions with a second cation. The R119A mutation may have altered the electrochemical gradient within the channel, thus requiring additional cations to support structural stability, and the introduction of the second cation reclaims structural integrity.

**Figure 4.**
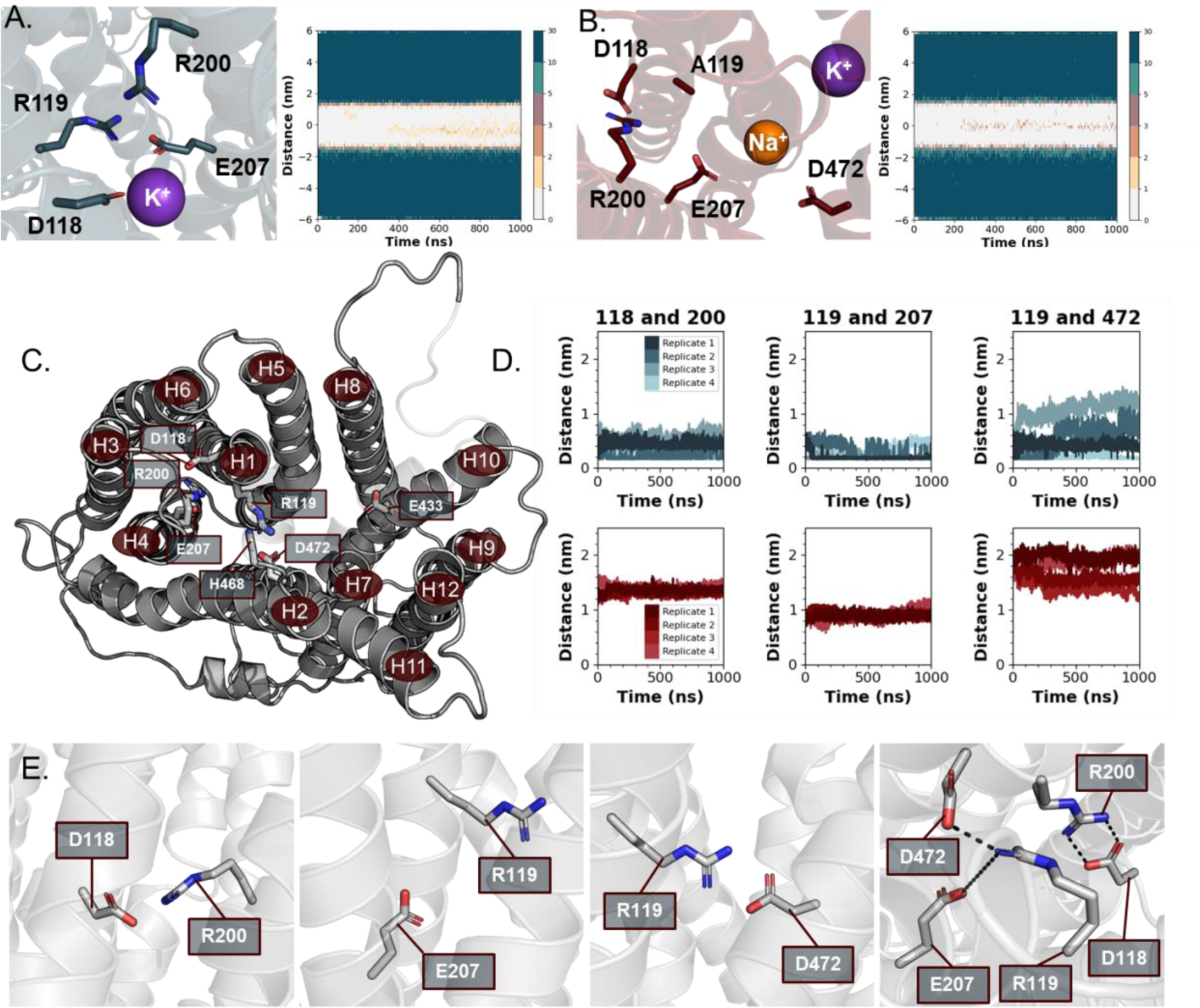
The R119A mutation disrupts a distinctive salt-bridge network within the Spns2 membrane-bound domain via alteration of the electrochemical gradient. A. Spns2_WT_ salt-bridge forming residues with ions within the channel. B. Spns2_R119A_ salt-bridge forming residues with ions within the channel. C. top-down visual of charged residues within the channel displayed, labeled with individual helices. D. minimum distance calculations of residues involved in salt-bridge network. E. Salt-bridge forming residue pairs.

Additionally, cation-membrane distance calculations over time revealed that the cation enters the channel and becomes oriented near the inter-bundle salt-bridge network (**Figure 4A-B**). This interaction could play an important role in salt-bridge breakage and formation, a process that occurs during conformational shifts from inward-facing to outward-facing during substrate translocation.(*15*) To observe the interaction pipeline for cation translocation, we analyzed the residue-ion interactions over time. We observed that potassium interacts with multiple charged residues and is likely initializing channel insertion from the extracellular region (**Table S1**). Additionally, potassium interacts with salt-bridge forming residues known to play a role in structural stability including Glu207, further supporting the role of secondary transport in translocation in Spns2 (**Figure 4A**). With respect to Spns2_R119A_, multiple cations were observed to enter the channel, including mixed cations (**Figure 4B**), a potentially essential process to support the altered electrochemical gradient within the channel, but also abolishing the substrate translocation capabilities.

MFS members are predominantly known to utilize protons to support transport. Previous research has hypothesized that Spns2 follows a proton/H^+^ secondary active transport model based off changing protonation of residues known to drive conformational changes. Here, it is possible we are observing ionic mimicry of proton-coupled transport. We do not observe a fully inward-facing conformation space in our simulations displayed in previous work. Still, this alteration of the electrochemical gradient within the Spns2 channel influence by the R119A mutation provides insight onto the importance of this residue on secondary active transport mechanisms and that this mutation impacts the electrochemical gradient, thus requiring additional protons/charged ions to re-stabilize. Observing how the ion-salt-bridge network relationship connects can further support our understanding of transport mechanisms.

MFS proteins are known to have interbundle salt-bridge networks oriented near the substrate binding domain (**Figure 4C**).(*15*) The salt-bridge networks have been observed in both outward-facing and inward-facing conformations and is important in structural conformation shifts. Literature has demonstrated Glu207 may protonate Arg119, two residues highly conserved in MFSs and known to be involved in the interbundle salt-bridge network, influencing substrate binding.(*39*) Additionally, Asp118 and Arg200 have been shown to be important salt-bridge forming residues, serving as a master switch with Glu207.(*38*) Analysis of the distances between these residues indicated that residues 119-207 and 118-200 interactions approach potential salt-bridge distances in Spns2_WT_ (average distance 0.25 ± 0.05 nm and 0.37 ± 0.04 nm, respectively) (**Figure 4D-E, S7**). Analysis of all potential salt-bridge forming residues were explored for their salt-bridge interactions, which revealed 468-472 interaction to be an additional salt-bridge forming complex in Spns2_WT_ (**Figure S7**).

When comparing Spns2_WT_ and Spns2_R119A_, we observe that there are some differences between the 119-207 and 118-200 interactions. Both pairs of residues are further away compared to Spns2_WT_ (119-207 average distances 0.90 ± 0.03 nm and 118-200 average distances 1.34 ± 0.02 nm). Spns2_R119A_ demonstrated differences in interactions between 119-207 and 118-200, but not the 468-472. Analysis of the ion-salt-bridge network interactions displayed that K^+^ interacts with Glu207, and a change in residue-residue interactions with Glu207 may impact the secondary active transport mechanism. These results indicate the formation of two salt-bridge interactions, 119-207 and 118-200, are essential for substrate translocation. R119A leads to a disruption within the salt-bridge network which has potential to alter the gating and cavity helices, impacting substrate translocation.

### Spns2 and S1P binding initialization

To determine which systems simulated with S1P had successful protein-ligand interactions, minimum distance calculations were performed per-residue to the center of S1P. For Spns2_WT_, only two of the six simulations constructed with S1P resulted in protein-ligand interactions (**Figure 5A, S8**). Out of those two (replicates 2 and 5), only one resulted with S1P entering the Spns2 channel, while the other resulted in S1P being lodged between Spns2 and the membrane (**Figure 5B**). While unexpected, analysis of the interactions between these two replicated can provide some preliminary understanding of the initial protein-ligand interactions that lead to translocation. None of the Spns2_R119A_ systems demonstrated consistent binding interactions with S1P (**Figure S9**).

**Figure 5.**
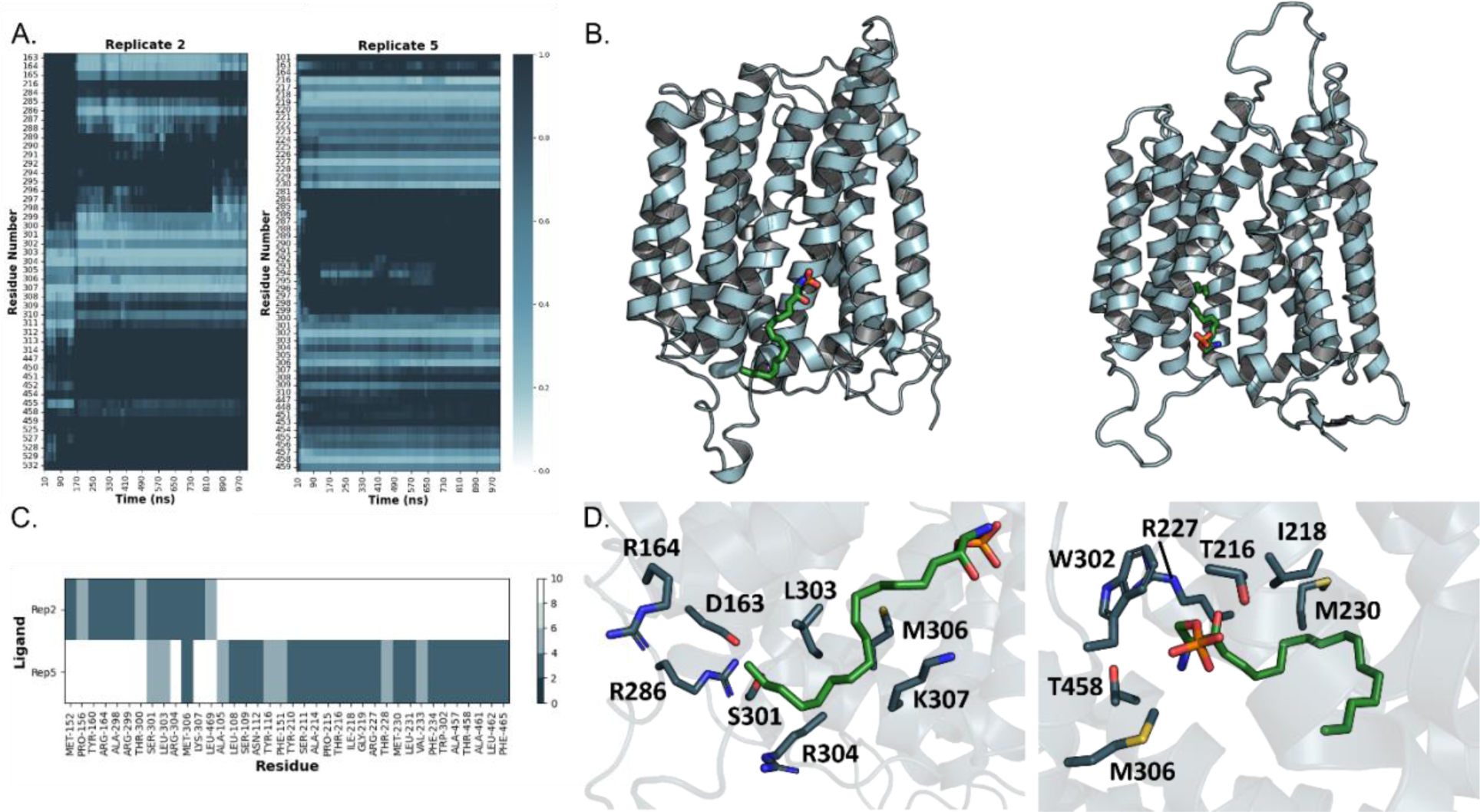
S1P initializes translocation via Spns2 by interacting with cytofacial residues 300-310. A. Minimum distance of individual residues to S1P over full simulation time. B. Structural view of S1P in Spns2 dominant morphologies. C. Protein-ligand interaction fingerprinting of replicate 2 and replicate 5. Fingerprinting was performed using an in-house python script. D. Visual representation of S1P interacting residues in the Spns2 dominant morphologies for replicate 2 (left) and replicate 5 (right).

Observing the minimum distance graphs with only features < 1 nm in distance extracted, both replicates initialize with residues 300-310 (**Figure 5A**). Replicate 2 shortly after shifts interactions with residues Asp163 and Arg164, not observed with replicate 5. Interestingly, this shift instigates interactions with Arg286, Ser301, Leu303, Arg304, Met306, and Lys307, mostly polar residues. Replicate 5 says consistent with the 300-310 residues, specifically interacting with Thr216, Ile218, Gly219, Arg227, Met230, Trp302, Met306, and Thr458. To further analyze the Spns2-S1P interactions, fingerprinting was performed extracting the minimum distance per residue over the last 200 ns of simulation (**Figure 5C-D**). The results support observations from the minimum distance analysis: replicate 2 interacting with Arg164, Ser301, Leu303, Arg304, Met306, and Lys307, while Rep5 interacting with Thr216, Ile218, Gly219, Arg227, Met230, Trp302, Met306, and Thr458. These results demonstrate that while initial interactions between Spns2 and S1P are driven by residues 300-310 residues, entrance into the Spns2 channel requires interactions with residues such as Thr216, Arg227, and Met230. The translocation pathway may require a more inward-facing morphology that the structure is unable to obtain due to protonation limitations in this study. This provides preliminary insight into S1P translocation via Spns2, and a route for drug discovery researchers to approach when developing novel Spns2 inhibitors.

Collectively, our observations suggest that Spns2 demonstrates a rocker-switch transport model. This transport mechanism is impacted by the R119A mutation, a residue involved in an inter-bundle salt-bridge network, suggesting that disruption to this network inhibits S1P translocation by altering secondary active transport and altering the electrochemical gradient within Spns2 channel. Further research should focus on identifying movement of S1P through the Spns2 channel to provide additional insight into how this process functions by utilizing steer-MD simulations or pre-docked S1P. These simulations should consider proton-mediated secondary active transport via altering system pH or by utilizing proton/H^+^ in simulation. Overall, this work shows insight into the initial Spns2 translocation mechanism, providing structural insight into drug discovery pathways for targeting S1P translocation for metastatic cancer.

### Conclusions

Spns2 has been shown as a promising drug target to control the metastatic microenvironment by suppressing S1P transport. Understanding the structural mechanism of Spns2 is essential for identifying the best avenues for drug development. Here, we utilize computational research approaches to deepen our understanding of the protein-structure function relationship and the impact of a single-site mutation, R119A, as it abolishes S1P transport. We identified that Spns2 follows a rocker-switch transport model and that a network of salt-bridge forming residue interactions is essential in substrate translocation. We revealed an inward-facing occluded state characterized by TM5 and TM11 occluding the cytofacial side of the channel, influencing potential S1P and inhibitor binding sites with implications for drug design. The Spns2_R119A_ system disrupts the inter-bundle salt-bridge network by altering the electrochemical gradient, thus abolishing S1P translocation. This disruption impacts both the gating helices, which is involved in inter-domain conformational shifts important in substrate translocation, and cavity helices, which are responsible for substrate binding within the channel. This details an attractive approach to reduce S1P transport by disrupting the salt-bridge network within the channel. This information provides new perspectives on how to target Spns2 for drug discovery purposes.

## Supplementary Figures

**Figure S1.**
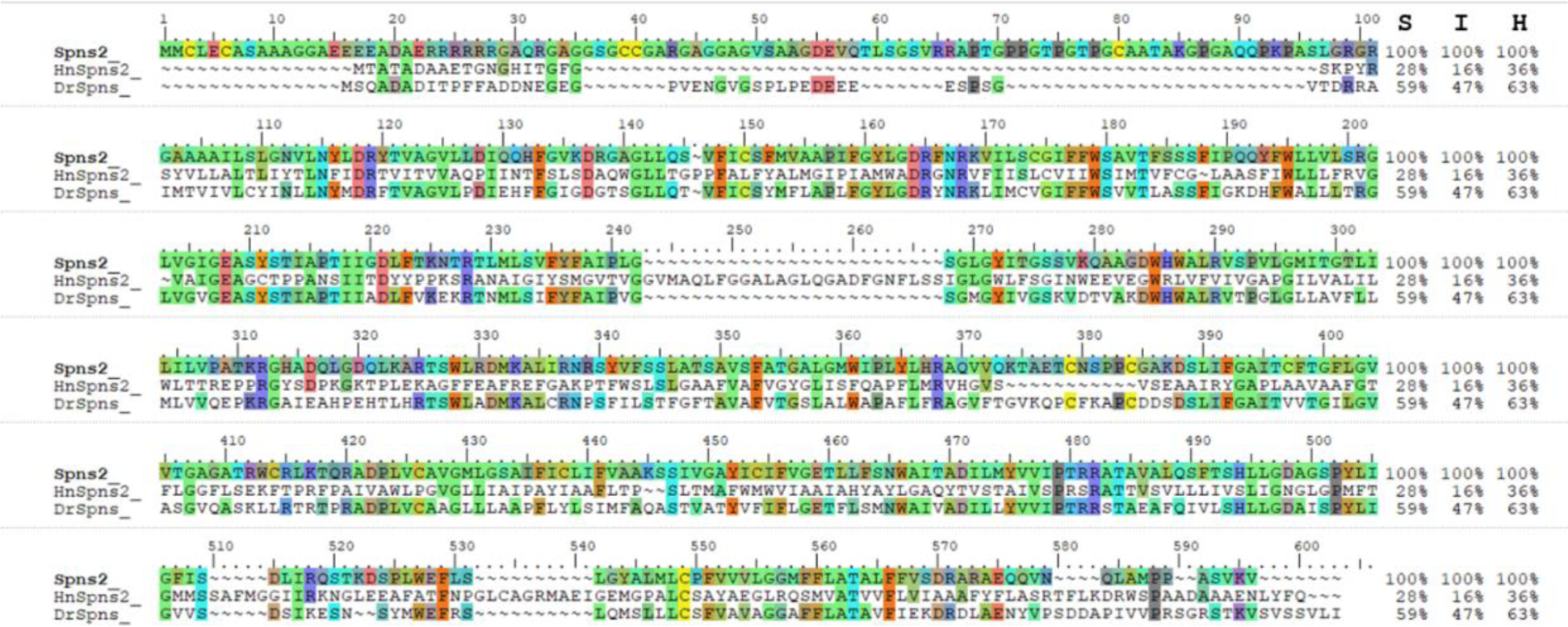
Sequence alignment of Spns2 and bacterial homolog Spns2. Sequence alignment was performed in Schrödinger-Maestro using human Spns2 (accession #Q8IVW8), *Hyphomonas neptunium* Spns2 (extracted from PDB 6e8j), and *Danio rerio*, or Zebrafish (accession #A2SWM2).

**Figure S2.**
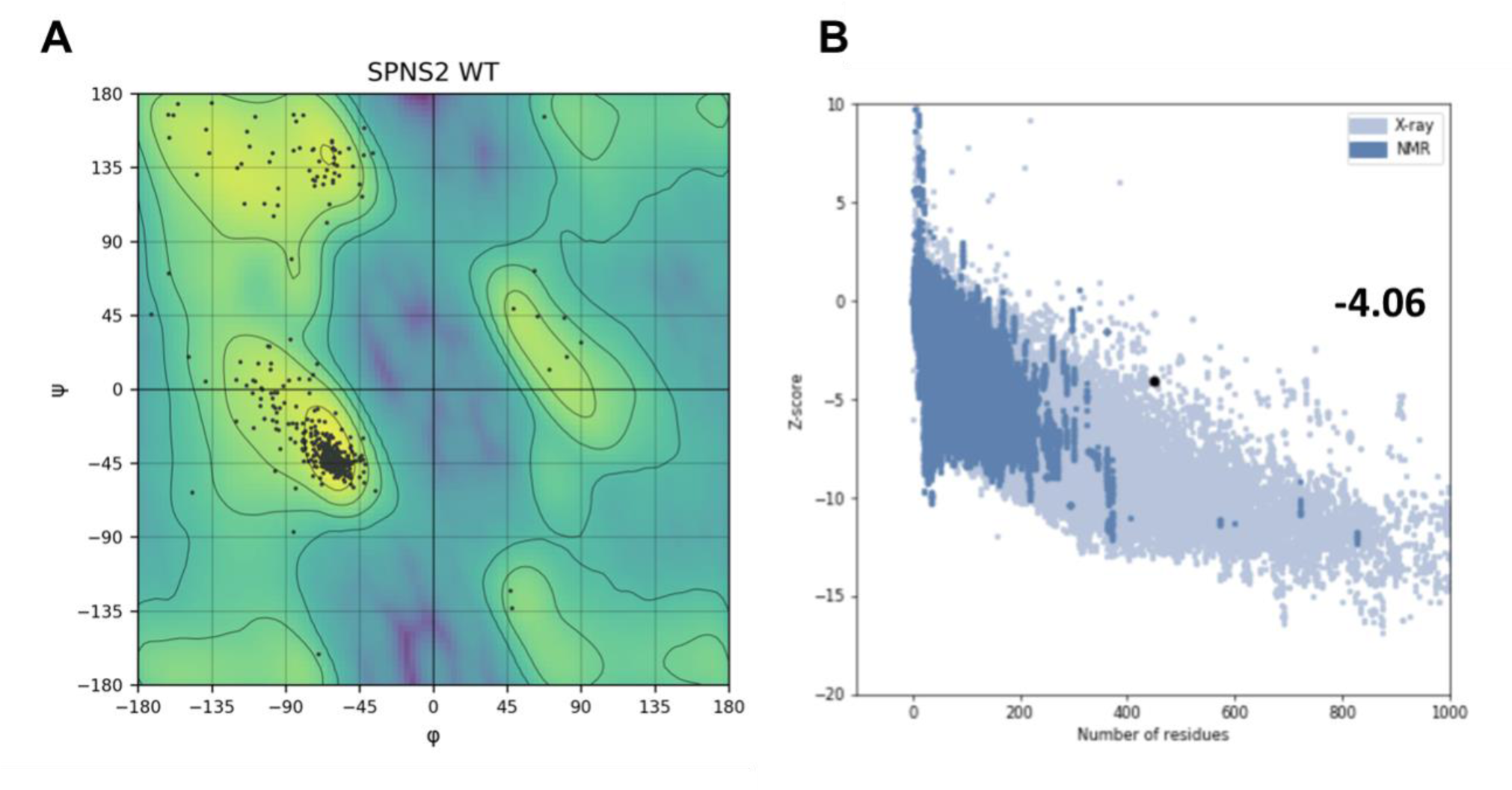
Spns2_WT_ homology model structure validation. A. Ramchandran plot analyzing the dihedral angles of the protein backbone. B. ProSA to score the model against known experimentally solved structures with a z-score of −4.06.

**Figure S3.**
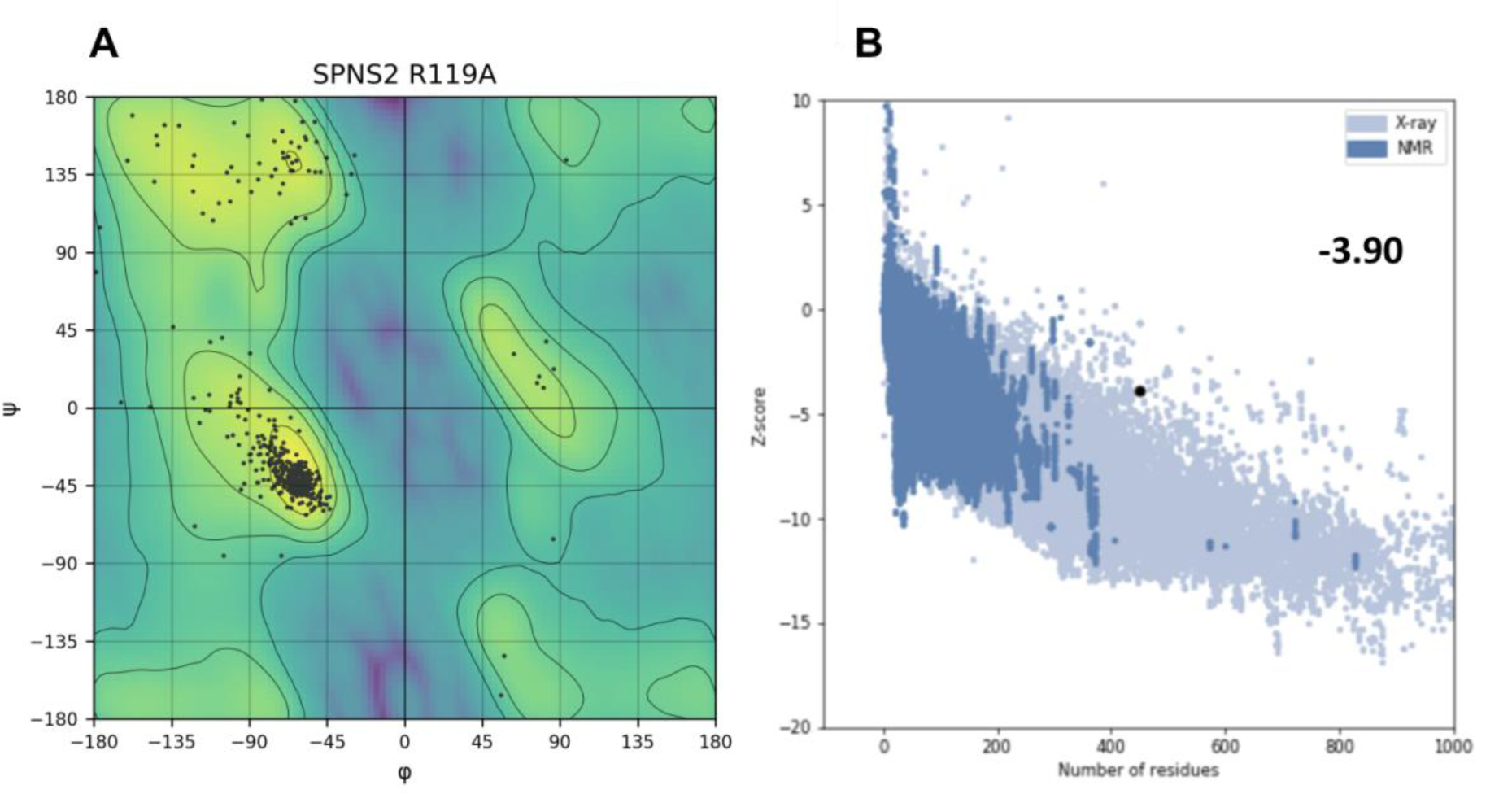
Spns2_R119A_ homology model structure validation. A. Ramchandran plot analyzing the dihedral angles of the protein backbone. B. ProSA to score the model against known experimentally solved structures with a z-score of −3.90.

**Figure S4.**
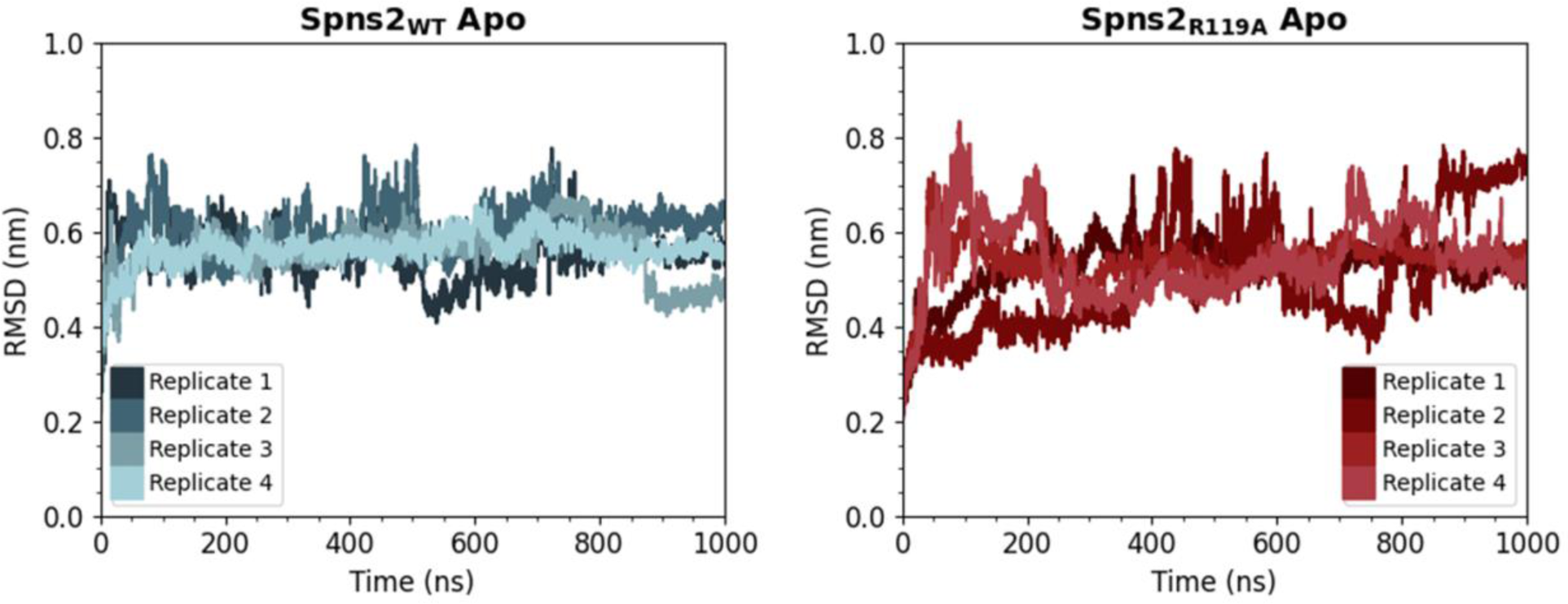
Root-mean square deviation of Spns2 Molecular Dynamics Simulations. RMSD was calculated using GROMACS rms feature with a 0.2-nm cutoff on individual replicates.

**Figure S5.**
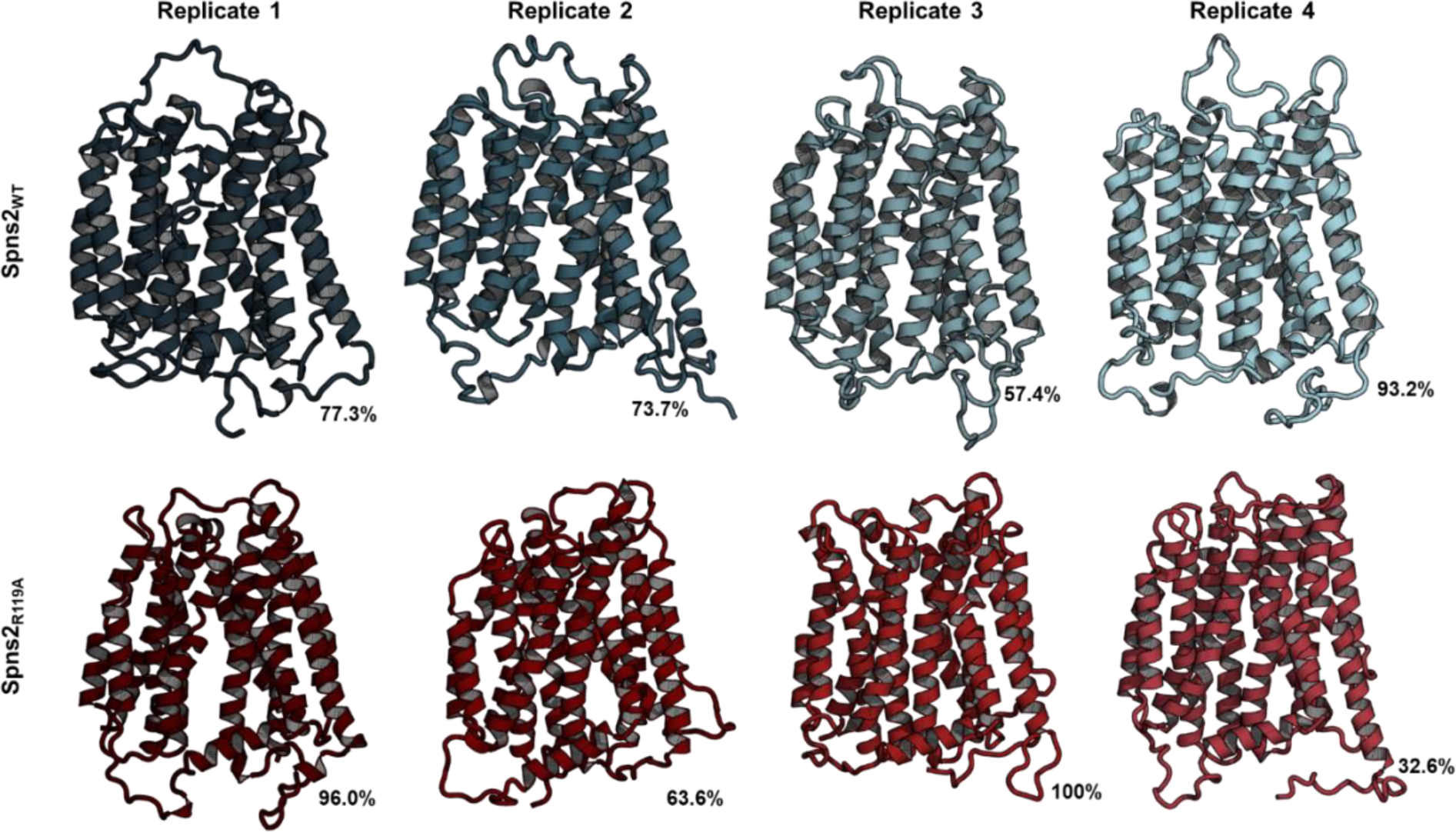
Per-replicate Molecular Dynamics dominant morphologies of apo systems. a. Spns2_WT_ clusters and b. Spns2_R119A_ clusters. Clustering was performed using GROMACS cluster feature using the gromos algorithm with a 0.2-nm cutoff.

**Figure S6.**
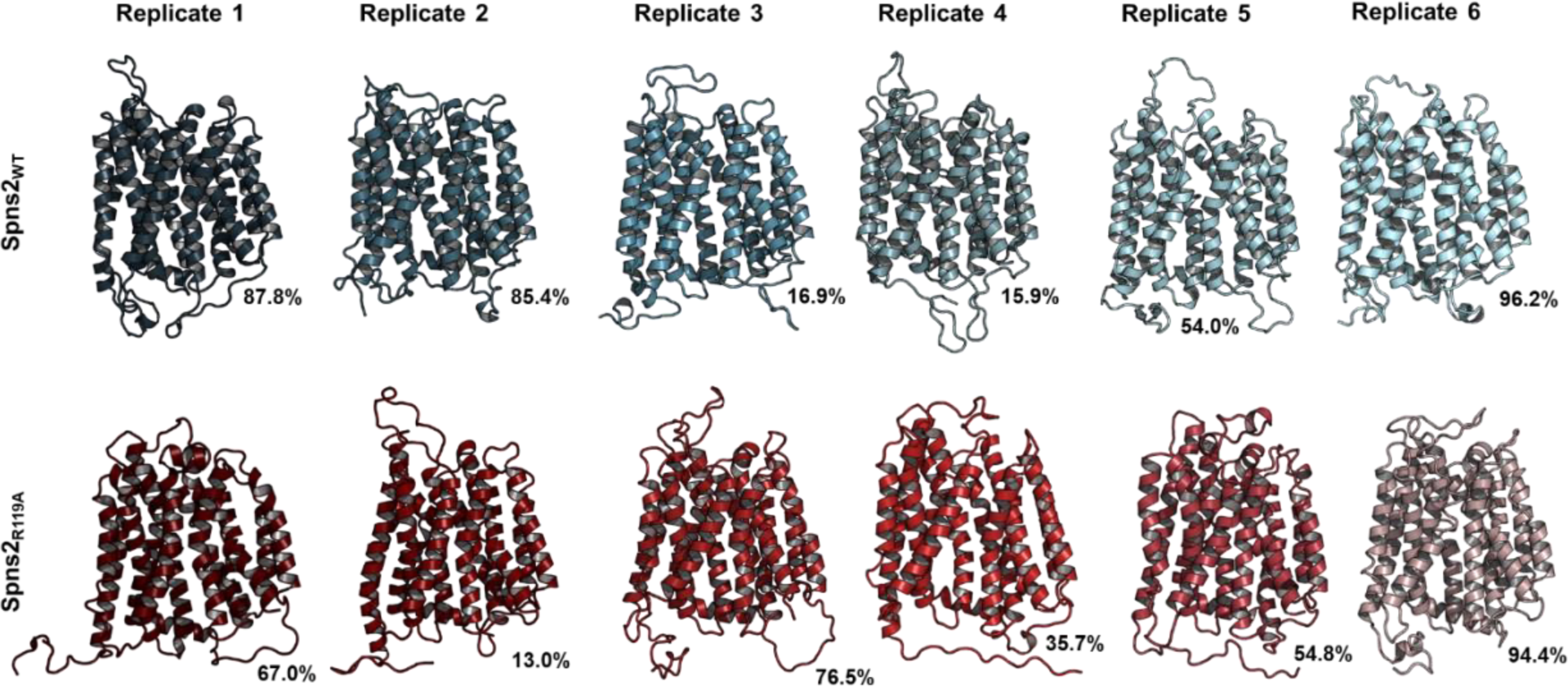
Per-replicate Molecular Dynamics dominant morphologies of S1P systems. a. Spns2_WT_ clusters and b. Spns2_R119A_ clusters. Clustering was performed using GROMACS cluster feature using the gromos algorithm with a 0.2-nm cutoff.

**Figure S7.**
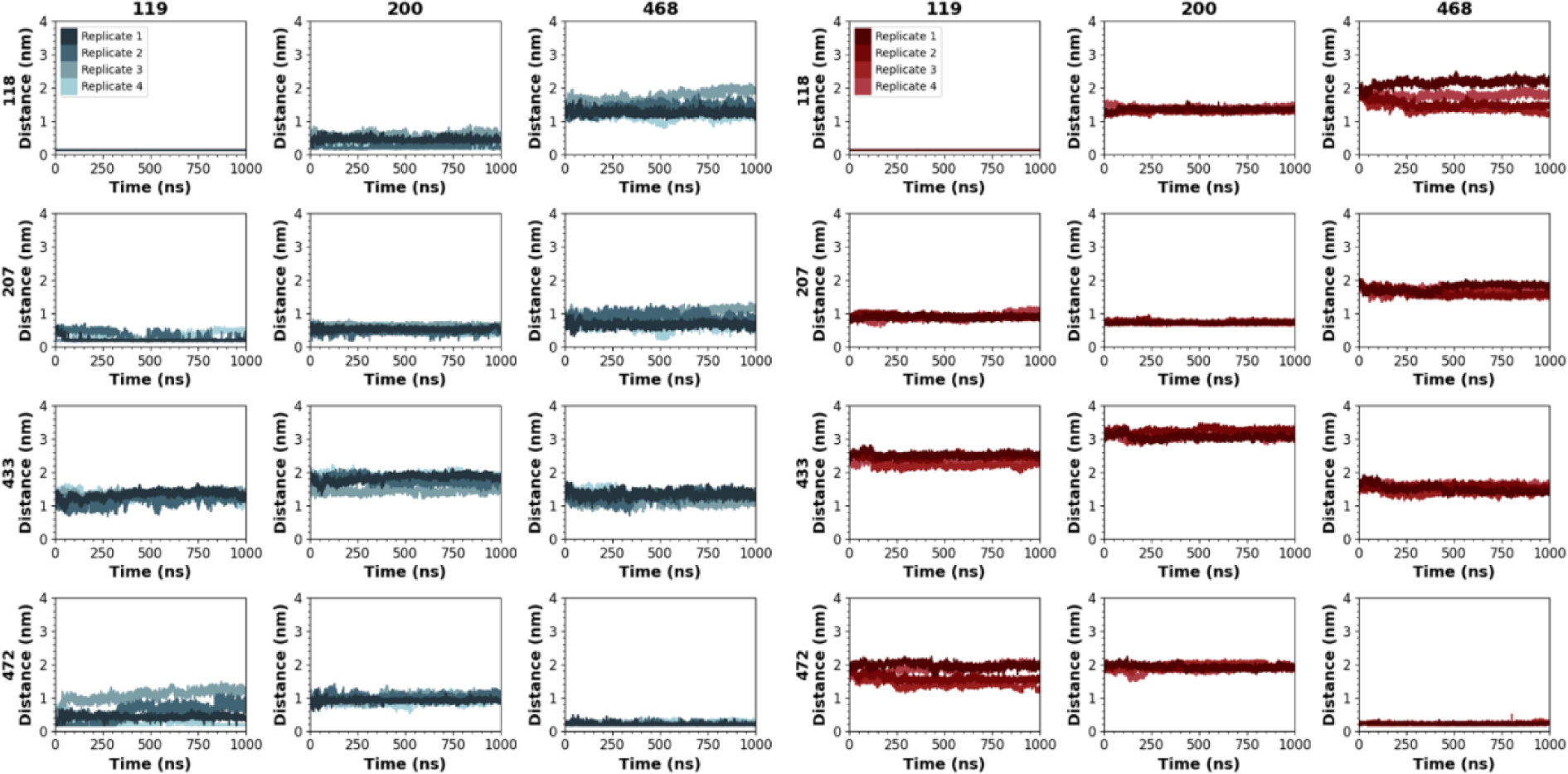
Distance calculations between potential salt-bridge forming residues within the membrane-bound domain of Spns2. Distance was calculated using the mindist feature in GROMACS. The blue colors represent residues within Spns2_WT_ and red represent Spns2_R119A_.

**Figure S8.**
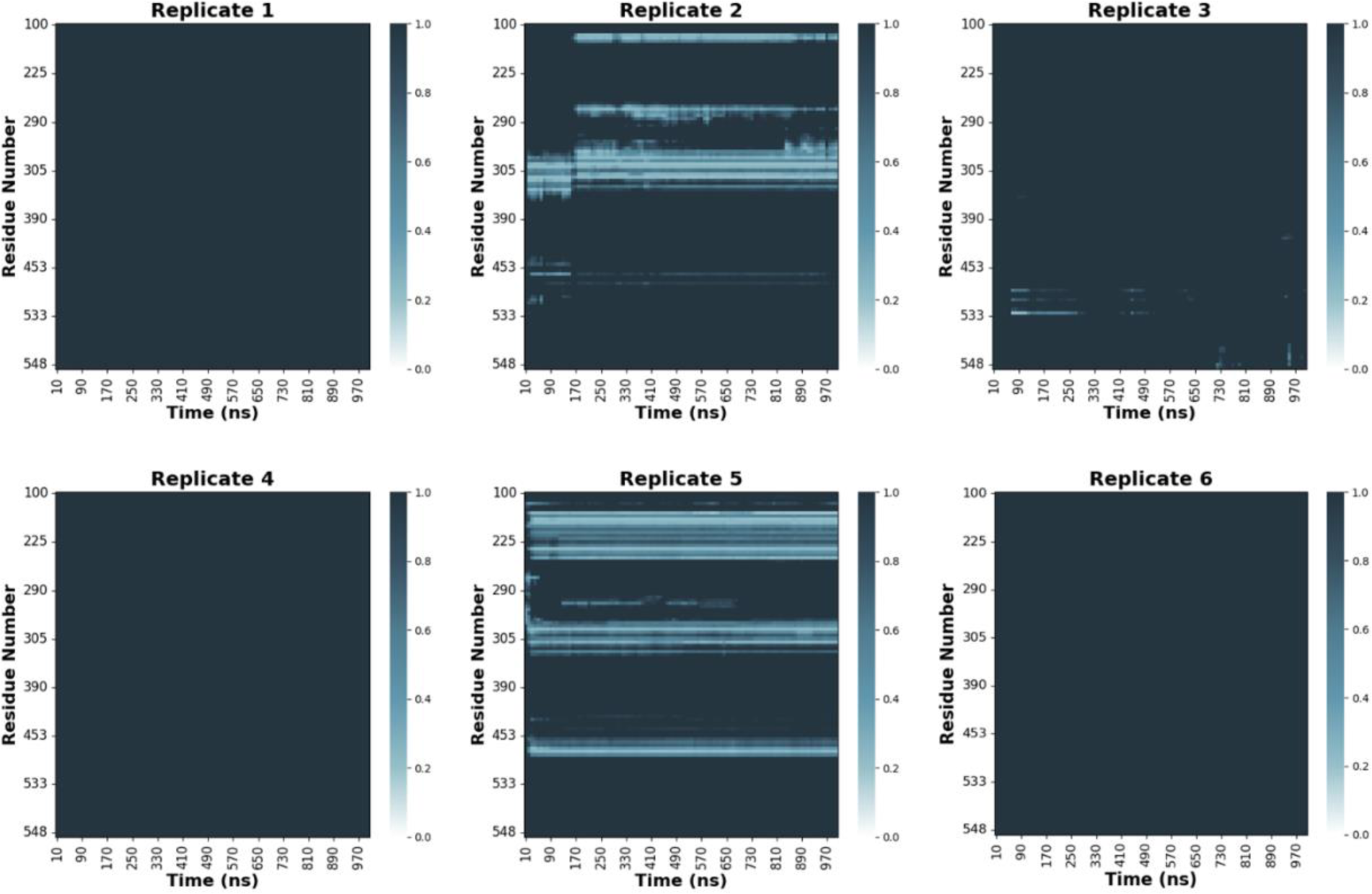
Minimum distance calculations between sphingosine 1 phosphate (S1P) and Spns2_WT_ residues. Minimum distance was calculated using the GROMACS mindist package. Distances above 1 nm are represented in deep blue, while distances < 1 nm are in lighter blue shades.

**Figure S9.**
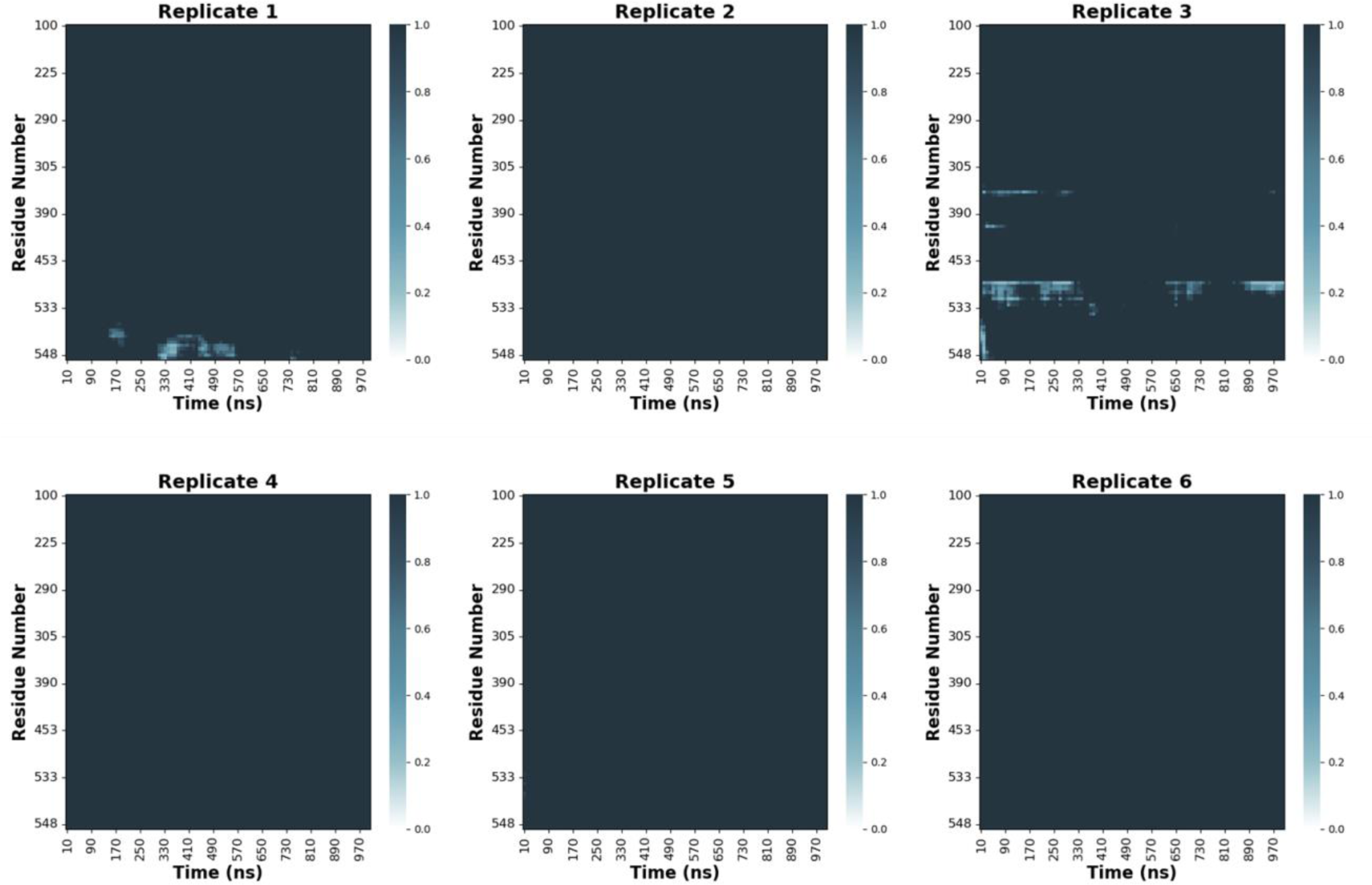
Minimum distance calculations between sphingosine 1 phosphate (S1P) and Spns2_R119A_ residues. Minimum distance was calculated using the GROMACS mindist package. Distances above 1 nm are represented in deep blue, while distances < 1 nm are in lighter blue shades.

**Table S1.** Residue-potassium interactions support a symport secondary active transport model.

| Extracellular Interactions |  | Channel Interactions | Cytofacial Interactions |  |  |
| --- | --- | --- | --- | --- | --- |
| Residue | Minimum Distance (Å) | Residue | Minimum Distance (Å) | Residue | Minimum Distance (Å) |
| Gln254 | 2.69 | Tyr116 | 4.52 | His288 | 2.83 |
| Arg487 | 2.71 | Glu207 | 4.64 | Gln291* | 2.85, 3.38 |
| Gln488 | 4.29 | Ser211 | 4.57 | Arg394 | 4.83 |
| Thr490 | 4.07 | Tyr235 | 3.83 | Tyr449 | 4.30 |
| Lys491 | 4.82 | Phe437 | 4.51 | Phe526 | 4.39 |
| Asp492 | 2.93 | Trp440 | 3.01 |  |  |
| Trp496 | 4.95 | Ser464 | 4.27 |  |  |
| Lys136 | 4.91 | Ser467 | 4.21 |  |  |
| Asp137 | 2.61 | His468 | 2.63 |  |  |
| Arg342 | 4.14 |  |  |  |  |
| Ser483 | 4.77 |  |  |  |  |
| Arg487 | 4.73 |  |  |  |  |
| Glu497 | 2.50 |  |  |  |  |
| Thr352 | 4.65 |  |  |  |  |

(\*) Interactions with multiple ions

